# Phosphorylation patterns of pre-ribosomal proteins associated with the RNA exosome

**DOI:** 10.64898/2026.01.29.702549

**Authors:** Felipe A. Almeida, Valdir Gomes Neto, Rebeca B. Jantsch, Moises R. A. Barros, L. Paola P. Cepeda, Bruno R. S. Queiroz, Anna B. Machado, Ana P.J. Menezes, Julia P. C. Da Cunha, Carla C. Oliveira

**Affiliations:** Department of Biochemistry, Institute of Chemistry, University of São Paulo, 05508-000, São Paulo, SP, Brazil; Laboratory of chemistry and function of proteins and peptides, Center for Biosciences and Biotechnology, State University of Northern Fluminense Darcy Ribeiro, 28013– 600, Campos dos Goytacazes, RJ, Brazil; Cell Cycle Laboratory, Center of Toxins, Immune Response and Cell Signaling-Center for Research on Toxins, Immune-response, and Cell Signaling (CeTICS), Butantan Institute, São Paulo 05503-900 SP, Brazil

## Abstract

The RNA exosome is an essential and ubiquitous RNase with exonucleolytic activity, involved in ribosome biogenesis and RNA quality control in eukaryotes. It is present both in nucleus and cytoplasm, and interacts with specific cofactors in each cell compartment, which are essential for recruitment and activity control of the exosome. Posttranslational modifications are known to regulate enzyme activity and protein interaction, although their precise roles are individually specific. In this study, we investigated the phosphorylation status of proteins associated with the nuclear (Rrp6) and core (Rrp46) subunits of the RNA exosome in *Saccharomyces cerevisiae*. Using co-immunoprecipitation followed by phosphopeptide enrichment and high-resolution mass spectrometry, we identified 121 phosphorylation sites on proteins functionally related to rRNA processing. Differential phosphorylation patterns between Rrp6 and Rrp46 co-immunoprecipitations are consistent with distinct exosome assemblies and suggest potential regulatory roles for phosphorylation. The results shown here highlight the role of phosphorylation in the recruitment and control of the exosome in RNA processing and degradation, offering new insights into the posttranscriptional control of gene expression.

## INTRODUCTION

The RNA exosome complex is a ubiquitous RNase with 3’-5’ exonucleolytic activity involved in processing and degradation of all classes of RNA in eukaryotes (Kilchert *et al*, 2016; Zinder & Lima, 2017). In budding yeast, the exosome is comprised of eleven different protein subunits, nine of which form the catalytically inactive core (Csl4, Rrp4, Rrp40-43, Rrp45, Rrp46, Mtr3). The core associates with Rrp44/Dis3 in the cytoplasm, forming Exo-10, and in the nucleus, Exo-11 is composed of Exo-10+Rrp6. Additionally, the exosome binds to specific cofactors in each subcellular compartment (Schneider & Tollervey, 2013). The exosome was first identified in the yeast *Saccharomyces cerevisiae* and later detected in other eukaryotes and some archaea species (Allmang *et al*, 1999b; Hartung & Hopfner, 2009; Mitchell *et al*, 1997). In the nucleus, the exosome plays an essential role in pre-rRNA processing and snoRNA and snRNA maturation (Allmang *et al*, 1999a).

The exosome is recruited to the 90S SSU processome, the first preribosomal particle to be formed cotranscriptionally, and is responsible for the degradation of the 5′-ETS spacer sequence after its release from the 35S pre-rRNA (Cepeda *et al*, 2019; Lau *et al*, 2021). Later during ribosomal maturation, the exosome is recruited to the pre-60S particles to process the 7S intermediate and form the mature 5.8S rRNA (Allmang *et al*., 1999a; Oliveira *et al*, 2002; Schuller *et al*, 2018; Thoms *et al*, 2015).

The major interactors of the exosome in the nucleus are responsible for its recruitment to pre-ribosomal particles during maturation and include the RNA helicase Mtr4, which is also a subunit of the TRAMP complex, and ribosome assembly factors Nop53 and Utp18, which recruit the exosome through Mtr4 (Bagatelli *et al*, 2021; Cepeda *et al*., 2019; Houseley *et al*, 2006; LaCava *et al*, 2005; Thoms *et al*., 2015). In the cytoplasm, the exosome is recruited by the SKI complex to degrade its substrates (van Hoof *et al*, 2000; Zhang *et al*, 2019). Ski7 is a critical adaptor connecting the SKI complex (Ski2-Ski3-Ski8) and the cytoplasmic exosome for mRNA degradation (Araki *et al*, 2001).

Although the structure and function of the exosome has been extensively studied in recent years, a detailed understanding of how the control of its function and recruitment is achieved is still lacking. Posttranslational modifications are a very important mechanism for controlling protein function, and protein phosphorylation is among the prevalent modifications. Phosphorylation of ribosomal proteins has long been known to influence translation (Bohlen *et al*, 2021; Cerezo *et al*, 2021; Nielsen *et al*, 1982), and has also been shown to affect ribosomal maturation and interaction between assembly factors (Ballesta *et al*, 1999; Filipek *et al*, 2024; Tomioka *et al*, 2018). Phosphorylation at specific sites on Dis3 (Ser809 and Tyr814) and Mrt4 (Thr1061 and Ser1067) from *Schizosaccharomyces pombe* has been shown to regulate exosome function (Telekawa *et al*, 2018). In this study, we investigated the role of phosphorylation in the regulation of the *Saccharomyces cerevisiae* exosome interaction with its cofactors. We selected Rrp6 and Rrp46 as representative components of the nuclear and core exosome complexes, respectively, to identify phosphorylation sites of proteins interacting with exosome-containing complexes that may be regulated during ribosome biogenesis, or cytoplasmic RNA processing/degradation processes.

## MATERIAL AND METHODS

### Yeast strains and culture conditions

Exosome subunits were affinity-purified from strains RRP46-TAP::HIS3 and RRP6-TAP::URA3 (Euroscarf) as described (Lourenco *et al*, 2013). Maintenance and growth of yeast cultures were performed according to standard procedures (Sherman et al., 1986). They were initially grown at 30°C in minimal medium (YNB) supplemented with 2% glucose, in addition to the essential amino acids, according to their auxotrophic marks. Yeast wild type strain W303 was used as negative control for subsequent analysis.

### Cloning and site-directed mutagenesis

For cloning of *RRP4* and *SKI7*, the plasmids were linearized by inverse PCR and the inserts were amplified from the yeast genomic DNA and recombinant plasmids were constructed with the InFusion HD (Clontech) recombination cloning system using competent Stellar (Clontech) cells. The site-directed mutagenesis was performed by overlap of PCR product and In-fusion DNA assembly kit, following the manufacturer’s instructions.

### Spot assay growth and growth curve

Precultures of the conditional strain *Δski7/yCP111-MET15::SKI7-TAP* (Neto *et al*, 2025) and *GAL1::13MYC-RRP4/yCP111-ADH1::RRP4-3HA*, expressing the corresponding plasmid-borne wild type Ski7 or Rrp4, or mutants (Ski7-S90A and Rrp4-S152A) in log phase were diluted for OD 600nm=0.1 and then subjected to 10-fold serial dilutions, and spotted onto glucose-containing solid minimal medium methionine-free or methionine-containing (2 mM) and incubated at 30°C. Growth curve measurements were performed in microplates incubated over 12h of growth using a BioTek LogPhase 600 Microbiology Reader set to read OD600 every 60 min at 30°C in the case of Ski7, and 30°C and 37°C in the case of Rrp4.

### Coimmunoprecipitation of Rrp46, Rrp6 and W303 strains

For the tandem affinity purification (TAP) method, these yeast strains were cultured in 6L YPD (1% yeast extract; 2% peptone; 2% glucose) medium. Biological triplicates were harvested at log phase (OD_600_ ∼ 1) by centrifugation at 8000 rpm (F12-6×500 LEX Rotor, Sorvall R6 C Plus) for 30 min at 4ºC. The cell pellets were resuspended in cooled extraction buffer (60 mM Tris–HCl pH 8.0, 50 mM NaCl, 10 mM MgCl_2_, 5% glycerol, 1 % PMSF, 1% triton, 1% EDTA, Pierce Protease Inhibitor Tablets, and Pierce Phosphatase Inhibitor Tablet) and flash-frozen in liquid N2.

The coimmunoprecipitation assays were carried out as previously described (Puig *et al*, 2001), with minor modifications. Briefly, the whole-cell lysates were obtained by cryogenic milling on a Ball Mill device (Retsch PM 100) and cleared by ultracentrifugation at 28,100 rpm (P50AT4 rotor, CP80NX Hitachi) for 60 min at 4°C. The lysates were incubated for 2 h at 4°C with 200 μl IgG-Sepharose 6 Fast Flow (GE Healthcare) previously equilibrated with extraction buffer and centrifuged at 500 rpm at 4ºC for 2 min. The beads were washed twice with extraction buffer and twice with wash buffer (50 mM tris-HCl and 50 mM NaCl). Elution was performed in the same column by adding 1 ml of TEV cleavage buffer and 100 units of TEV protease. The beads are rotated overnight at 4ºC and the eluate was recovered by gravity flow. Eluates were lyophilized and stored at -80ºC until treatment for LC–MS/MS analysis. Both input and eluate fractions were separated for analysis by SDS-PAGE and western blot.

### SDS-PAGE and immunoblot

Aliquots (∼5%) of yeast coimmunoprecipitated proteins were resuspended in SDS sample buffer, resolved on 10% SDS-PAGE (Tris-glycine running buffer), transferred to nitrocellulose membranes (Ambion) and incubated with antibody against the CBP tag (Millipore). Secondary antibody conjugated to IRDye680RD (anti-rabbit IgG, LI-COR) was employed, and near-infrared Western blot detection was carried out using ChemiDoc MP Imaging System (BioRad).

### Peptide digestion and TiO_2_ phosphopeptide enrichment

The lyophilized eluates were subjected to in-solution digestion for LC– MS/MS analysis. Briefly, proteins were resuspended in 50 mM NH_4_HCO_3_, 8 M urea and reduced with DTT at a final concentration of 5 mM at 56°C for 25 min. Samples were then alkylated with 14 mM iodoacetamide for 30 min under protection from light. After dilution with 50 mM NH_4_HCO_3_, samples were digested for 16 h with sequencing-grade modified trypsin (Promega) at a 1:50 (E:S) ratio at 37°C. The proteolysis was stopped by adding TFA to a final concentration of 0.4% (v/v) and the resultant peptides were desalted using SepPak tC18 cartridges (Waters).

A High-Select TiO_2_ kit (Thermo Fisher Scientific) was used to enrich phosphopeptides from tryptic peptides. The digested peptides were placed into TiO_2_ spin tips and the enrichment steps were performed according to the manufacturer’s instructions. After elution of the phosphopeptides, the eluate was vacuum-dried and stored at -80 ºC until mass spectrometry analysis.

### Mass spectrometry analysis and data acquisition

Digested peptides were resuspended in 0.1% formic acid (FA) and separated on an in-house reversed-phase capillary emitter column (10 cm × 75 μm, filled with 5 μm particle diameter C18 Aqua resins-Phenomenex) coupled to a nano-HPLC (Thermo). Mobile phases consisted of 0.1% FA in water (buffer A) and 0.1% FA in acetonitrile (buffer B). Peptides were eluted at 300 nL min^−1^ using a 60 min gradient (2–28% B for 45 min, 28–80% B for 13 min, and a return to 5% B in 2 min). The eluted peptides were analyzed by an LTQ-Orbitrap Velos (Thermo Scientific) (source voltage of 1.9 kV, capillary temperature of 200°C). The mass spectrometer was operated in DDA mode with dynamic exclusion enabled (exclusion duration of 45 seconds), MS1 resolution of 30,000, and MS2 normalized collision energy of 30. For each cycle, one full MS1 scan range of 200–2000 m/z was followed by ten MS2 scans (for the most intense ions) using an isolation window size of 1.0 m/z and collision-induced dissociation (CID). The 10 most intense precursor ions were selected for fragmentation by CID in DDA. The .raw data from the LTQ Velos-Orbitrap were analyzed with the MaxQuant (version 2.6.7.0) software using the UniProt database for Saccharomyces cerevisiae (strain ATCC 204508 / S288c), setting tolerance for the MS of 20 ppm and the MS/MS of 0.5 Da; carbamidomethylation of cysteine as a fixed modification; and phosphorylation at STY residues as variable modifications; and 1% False Discovery Rate (FDR). The match between runs option was used to increase the number of trusted IDs. The normalized Intensity values of the Phospho (STY)Sites.txt output and the Intensity columns for each biological replicate were used for downstream analyses. Identifier for the raw mass spectrometry data: PXD073587.

### Label-free phosphoproteomics data analysis

Initial data analysis was performed using Perseus software (version 2.1.3.0, Max Planck Institute of Biochemistry). The affinity purification assays using different baits were analyzed together. The MaxQuant output data was initially filtered removing contaminants, hits to the reverse database, and phosphoproteins with localization probability ≥ 0.70. The intensities were log2-transformed and the biological triplicates were grouped by categorical annotation in the two conditions assessed, Rrp46 and Rrp6. Furthermore, phosphopeptides were filtered to include only those detected in at least two of the three biological replicates or present uniquely in one condition (presence/absence filtering for exclusive identifications).

For differential phosphorylation analysis, intensity values log_2_-transformed were subjected to Student’s t-test. Only phosphopeptides with statistically significant differences (FDR-adjusted p-value (Benjamini–Hochberg) ≤ 0.05) were considered for downstream analysis. Additionally, phosphopeptides identified in control W303 samples were excluded to reduce false-positive identifications.

## RESULTS

To investigate the possible regulatory role of phosphorylation in the recruitment of the exosome to pre-ribosomal complexes, we conducted a label-free phosphoproteomic analysis using *Saccharomyces cerevisiae* strains expressing TAP-tagged versions of the nuclear (Rrp6) or core (cytoplasmic/nuclear - Rrp46) subunits of the RNA exosome. These subunits interact with distinct protein partners and ribonucleoprotein complexes during ribosome biogenesis, or cytoplasmic RNA degradation, and our goal was to identify phosphorylation events that may modulate these interactions. Protein co-immunoprecipitation experiments were therefore performed using total extracts of yeast cells expressing either Rrp6-TAP or Rrp46-TAP. Eluted proteins from biological triplicates were then digested with trypsin and resulting peptides were subjected to phosphopeptide enrichment and identification by mass spectrometry. The protein baits were detected both by mass spec and immunoblots (Fig. 1A). Extracts from wild-type W303 strain, processed in parallel, were used and proteins/phosphosites detected in control runs were excluded from downstream analyses to reduce background binders. The similarity between replicates was compared by a principal component analysis (PCA), which show two clusters, one for each sample, Rrp46 or Rrp6, with good representation (Fig. 1B).

**Figure 1.**
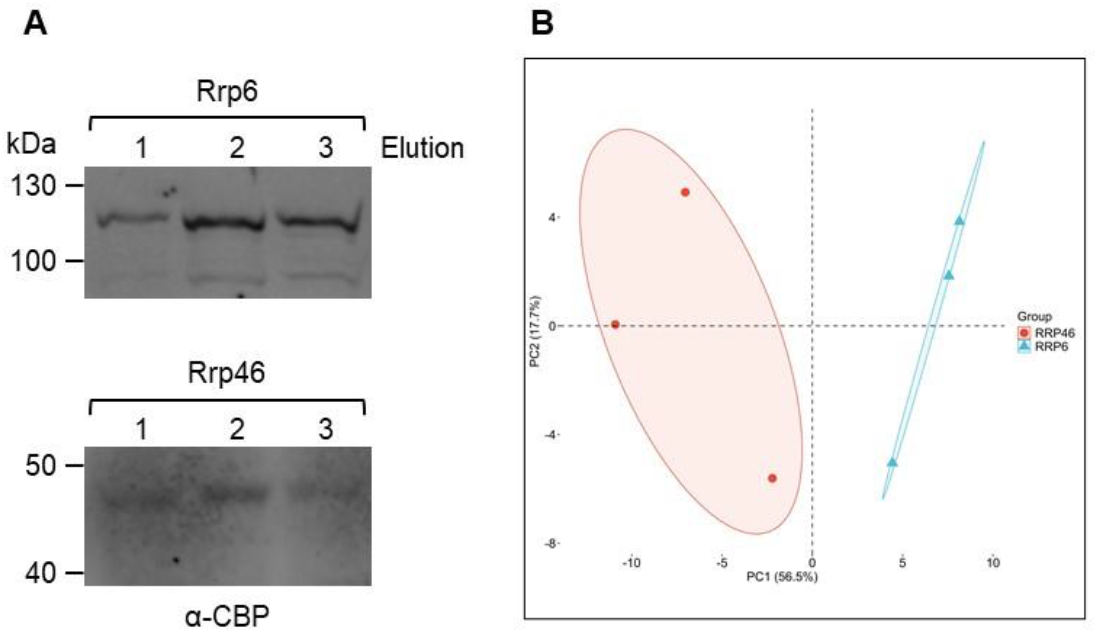
Protein co-immunoprecipitation with Rrp6-TAP or Rrp46-TAP. **(A)** Western blot validation of TAP-tagged RRP6 and RRP46 used as baits in co-immunoprecipitation assays. The immunoblot shows the bands corresponding to the baits Rrp6-TAP (∼105 kDa) and Rrp46-TAP (∼45 kDa) from the elution franctions, confirming efficient binding to the column. The TAP tag (∼21 kDa) accounts for the shift from native molecular weights (∼84 and ∼25 kDa, respectively). (B) Principal component analysis (PCA) of phosphopeptide intensity profiles. Each point represents one biological replicate of Rrp6 (blue triangles) or Rrp46 (red circles). PC1 and PC2 account for 56.5% and 17.7% of the total variance, respectively. Ellipses represent 95% confidence intervals for each group. Tight clustering of Rrp6 replicates and broader dispersion of Rrp46 samples indicate group specific phosphoproteomic profiles and differential variance structure.

A total of 756 phosphorylated peptides were identified (Suppl. Table 1), corresponding to phosphosites present on proteins co-purifying with the exosome subunits. Gene ontology analysis of proteins identified in these affinity-purified fractions showed prevalence of rRNA processing factors, which was expected due to the essential role played by the exosome in the ribosome maturation pathway (Fig. 2). Among the rRNA processing factors recovered, those involved in 5.8S rRNA maturation were the most abundant (Fig. 2), which reinforces the validation of our results, since the exosome is directly involved in mature 5.8S formation. From the initial peptides recovered, we selected a total of 121 phosphorylated residues that correspond to proteins functionally related to RNA processing for further analyses.

**Table 1.**
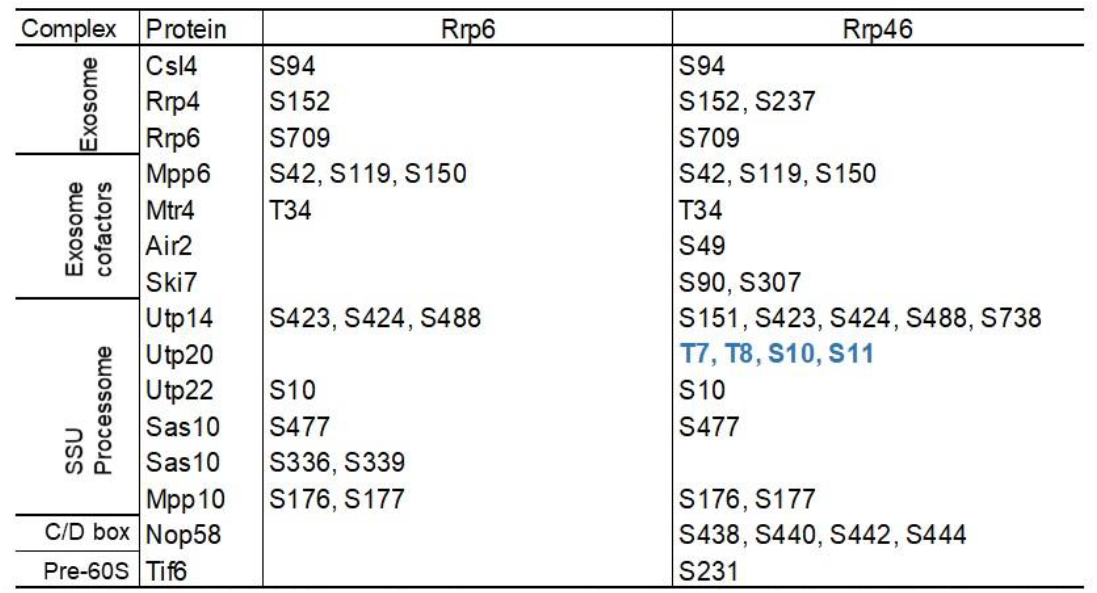
Phosphosites in identified peptides from proteins co-purified with Rrp6-TAP or Rrp46-TAP. Partial list of proteins identified, highlighting exosome subunits and ribosome maturation factors, and indication of phosphoresidues in each protein. Novel phosphosites are highlighted in blue.

**Figure 2.**
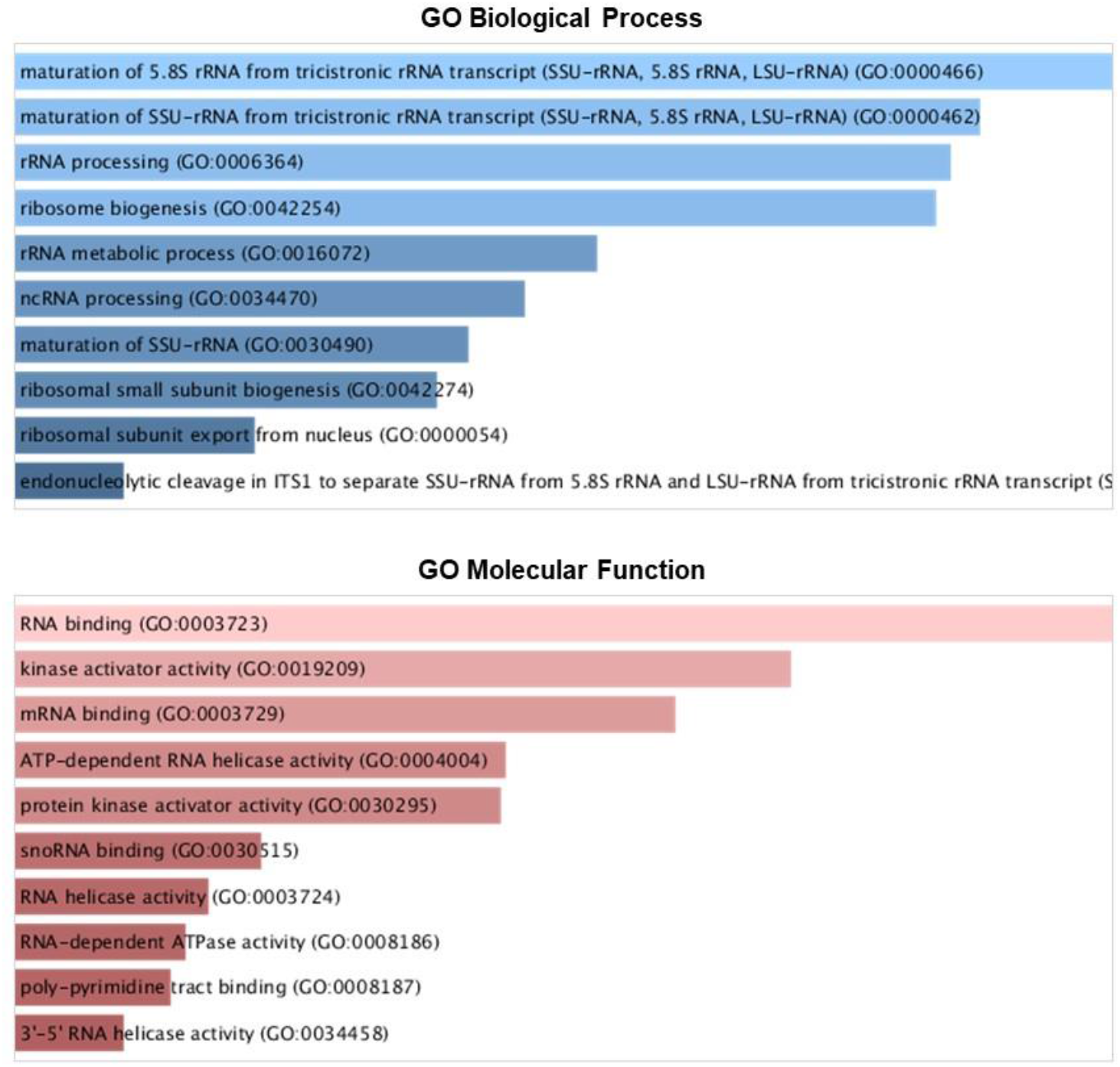
GO terms of biological processes and molecular function of the proteins co purified with the exosome subunits Rrp6 and Rrp46. **(A)** Biological process: Ribosome biogenesis factors were most abundant category in the samples. (**B**) Molecular function: RNA binding was the prevalent category. Three biological replicates were used in the purifications for the bait proteins were performed.

We investigated the distribution and intensity of phosphorylation events across the two experimental conditions. A heatmap based on phosphosite intensities for rRNA processing-associated proteins revealed both shared phosphorylation profiles and condition-associated patterns (Fig. 3).

**Figure 3.**
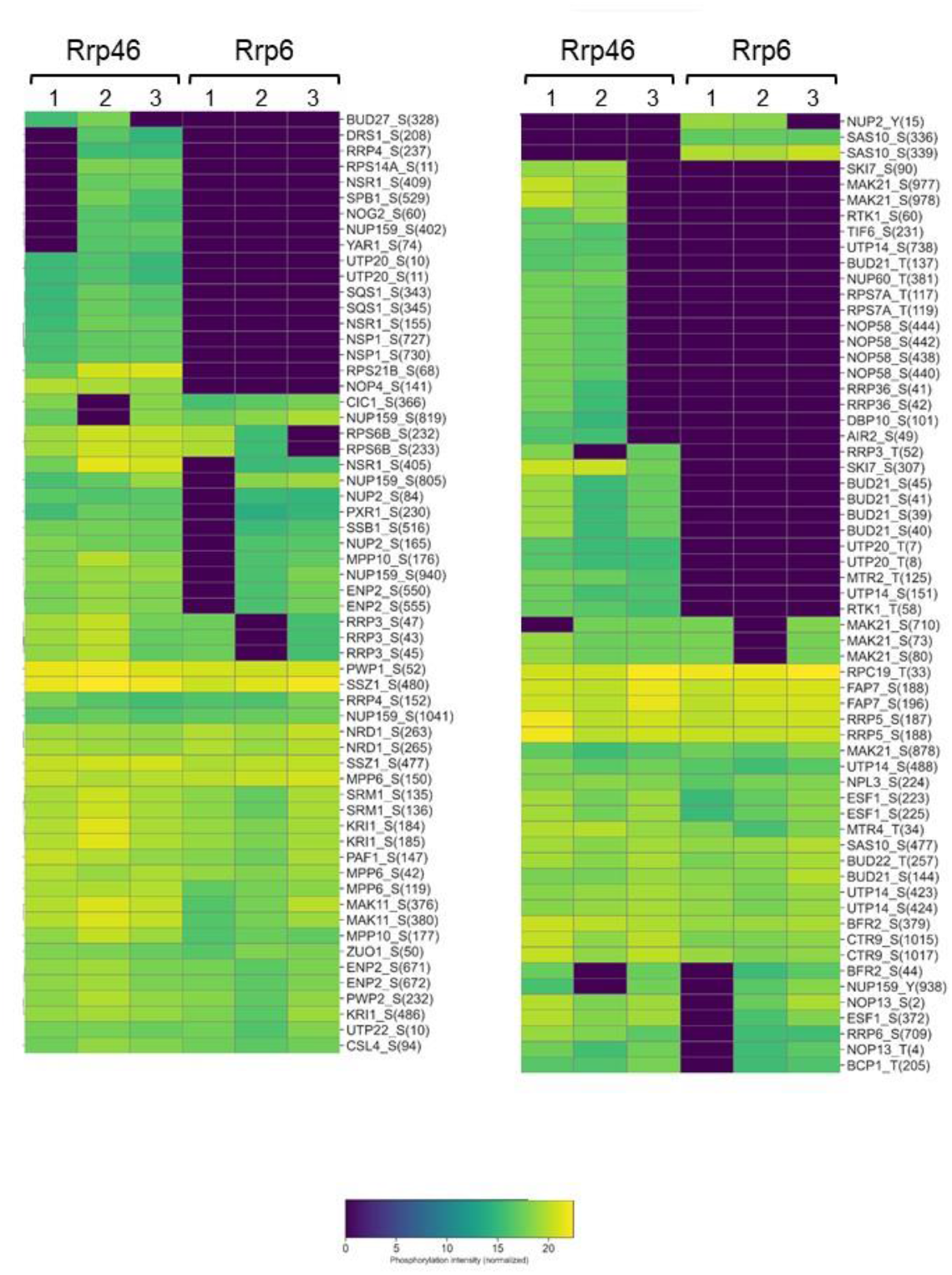
Heatmap of normalized phosphorylation intensities for rRNA processing-associated residues across Rrp6 and Rrp46 immunoprecipitates. Color gradients reflect relative phosphorylation levels across three biological replicates. Distinct clusters represent residues exclusive to Rrp6, Rrp46, or shared between both, highlighting phosphorylation-dependent specificity in exosome-associated preribosomal complexes. Numbers indicate the individual biological replicates used in the co-ip experiments.

Utp20, Nop4, Sqs1, and Nog2, among other ribosome maturation factors, were uniquely detected in Rrp46 samples, whereas other assembly factors, including Enp2, Pwp2, Kri1 and Utp22 were phosphorylated in both samples (Fig. 3), suggesting a heterogeneity of nuclear exosome complexes. Notably, the phosphosites on the N-terminal unstructured portion of Utp20 detected here have not been previously reported. The selective enrichment of phosphorylated peptides in Rrp46 samples suggests that the association of the exosome at different stages of ribosomal maturation may be mediated by additional assembly factors, other than Utp18 (SSU processome) and Nop53 (pre-60S) via Mtr4-Rrp6 direct interaction (Schuch *et al*, 2014).

A smaller set of phosphosites was uniquely detected in Rrp6 samples. Among those, Sas10 (residues S336 and S339) and nuclear pore-associated export factor Nup2 (Y15) were consistently phosphorylated only in the Rrp6-associated fractions (Fig. 3). Sas10 is functionally linked to the SSU processome, and has been shown to bind Rrp6-Rrp47 heterodimer (Mitchell, 2010), confirming the interaction of the exosome with factors involved in early rRNA processing steps in the nucleolus (Cepeda *et al*., 2019). Further demonstrating the consistency of our results, proteins with multiple phosphorylation sites showed particular phosphorylation states, depending on the residue and the complex they were purified with. Utp20, for example, had its four phosphorylated serine residues purified exclusively with Rrp46, while Sas10-S336 and S339 were phosphorylated exclusively in Rrp6 samples. Sas10-S477 and Mpp10, on the other hand, a protein of the same complex as Sas10, were phosphorylated in both samples (Fig. 3; Table 1). These differential phosphorylation states might indicate regulation of protein function, affecting interactions throughout ribosomal maturation.

Several phosphosites exhibited comparable phosphorylation levels in both Rrp6 and Rrp46 complexes, suggesting that these modifications play constitutive regulatory roles irrespective of the context of the ribosomal maturation stage with which the exosome is associated. These shared sites were found in central assembly factors such as Mpp10, Utp22, Utp14, Rrp5, Kri1, Enp2, Bud21, Bud22, Esf1, and Pwp2 (Fig. 3). These proteins are essential for 90S pre-ribosome formation, and the conserved presence of their phosphorylated forms in both conditions indicates that their regulation is fundamental and that the Exo11 complex remains bound to pre-ribosomal particles at different maturation stages. Strengthening this hypothesis, Pwp1, which is also phosphorylated in both conditions, is an example of a protein identified here that participates in pre-60S maturation (Fig. 3).

In addition to the phosphorylated peptides co-purified with the exosome subunits, phosphosites of three different exosome subunits were identified, Csl4-S94, Rrp6-S709, and Rrp4-S152, S237 (Fig. 3; Table 1). Interestingly, these residues were phosphorylated in both samples, purified with Rrp6 and Rrp46, with the exception of Rrp4-S237, which was co-purified exclusively with Rrp46, suggesting that the phosphorylation of this residue is involved in cytoplasmic function of the exosome, either for activity control, or recruitment of the complex by its cofactors. Importantly, this residue is exposed on the exosome complex structure (Makino & Conti, 2013) (Fig. 4; Suppl. Fig. 1), indicating its suitability for being modified. Csl4-S94 and Rrp4-S152 phosphorylation have already been described (Synowsky *et al*, 2006), confirming the high reproducibility of our purification method. Csl4-S94 is present in a predicted disordered domain of this protein. Rrp4-S152, on the other hand, is imbedded in its S1 RNA binding domain and is accessible for post-translational modifications (Fig. 4; Suppl. Fig. 1), and is conserved in the human ortholog EXOSC2 (Ramos *et al*, 2006; Synowsky *et al*., 2006). The importance of Ser152 phosphorylation for Rrp4 function was assessed by substituting Ala for this Ser (Rrp4-S152A) and analyzing growth of yeast cells expressing this mutant. For this analysis, the endogenous promoter of *RRP4* was replaced by *GAL1* promoter, which is activated in the presence of galactose, and inhibited in glucose medium, and this strain (*GAL1::Myc-RRP4*) was transformed with a plasmid expressing either wild type Rrp4, or the mutant Rrp4-S152A, fused to the HA tag and under control of the constitutive promoter *ADH1*. Growth was then analyzed in glucose medium, condition in which only the plasmid-encoded Rrp4-HA proteins are expressed. Despite the evolutionary conservation of this residue, however, its change to the non-phophorylatable Ala does not affect growth (Fig. 5), suggesting that phosphorylation at position 152 does not interfere with the RNA binding activity of Rrp4.

**Figure 4.**
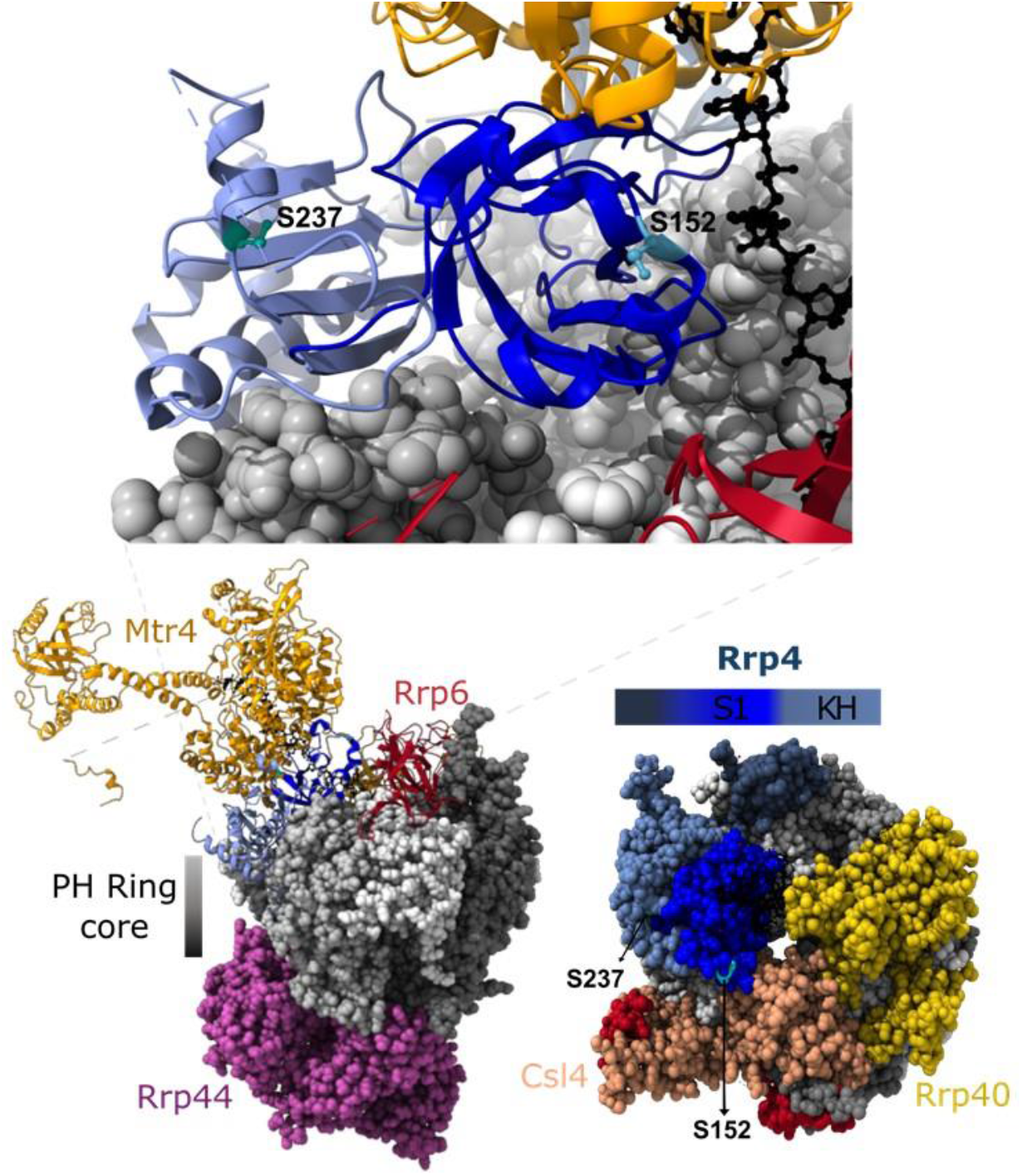
Structure of the exosome, highlighting the positions of Rrp4 phosphorylated residues Ser152 and Ser237 (indicated, cyan and green, respectively). Structure obtained from PDB 6FSZ.

**Figure 5.**
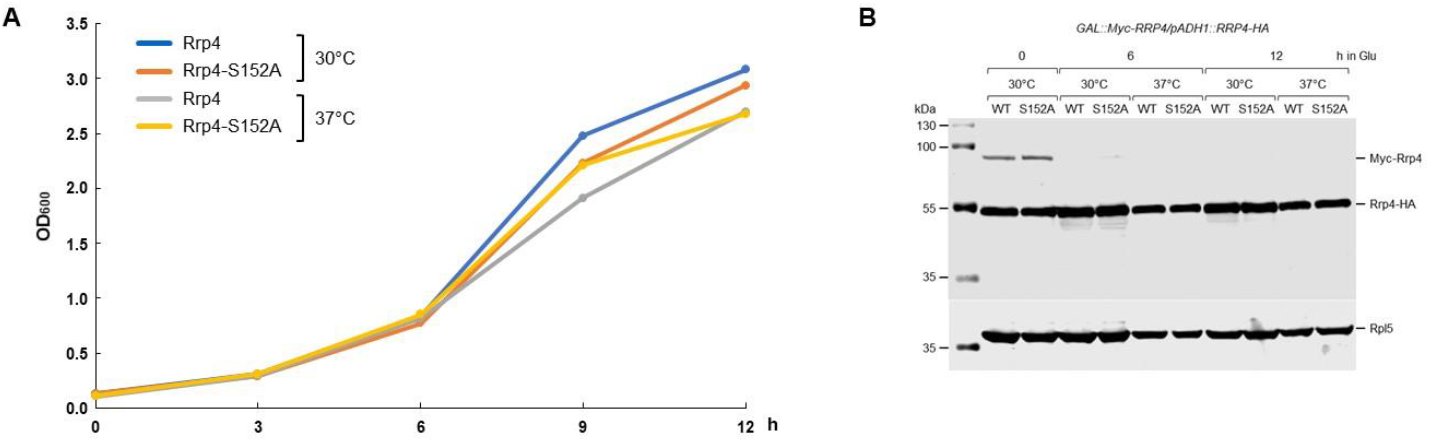
Effects of Rrp4-S152A mutation on cell growth. Plasmid-borne wild type Rrp4-HA: or mutant S152A were expressed in *GAL::RRP4-Myc* cells and growth was monitored over time in glucose medium, condition in which only the plasmid copies of Rrp4 are expressed. (**A**) Growth curve in glucose medium. (**B**) Western blot for detection of chromosomal-encoded Myc-Rrp4 and plasm id-encoded WT and mutant (S152A) Rrp4 fused to HA tag. Endogenous Rpl5 was used as a loading control.

Rrp6-S709 is present in the lasso domain, involved in RNA binding and stimulation of exosome activity, and is inserted in the canonical nuclear localization signal (NLS) of Rrp6 (697-723) (Gonzales-Zubiate *et al*, 2017; Wasmuth & Lima, 2017), potentially affecting both RNA binding and Rrp6 interaction with karyopherins. Phosphosites S42, S119, S150 of the exosome cofactor Mpp6 (Schilders *et al*, 2005; Wasmuth *et al*, 2017) were found in both samples (Table 1). Mpp6 has already been identified as a phosphorylated protein in high-throughput analyses (Leutert *et al*, 2023), validating our results. A minimal portion of Mpp6 comprising amino acid positions 90 through 118 was crystallized in complex with the exosome and shown to bind directly Rrp40 (Wasmuth *et al*., 2017). Due to its proximity to the Mpp6 portion binding the exosome, it is possible, therefore, that phosphorylation of S119 might influence the Mpp6 association with the exosome, possibly positively, since this phosphosite was recovered with both baits.

Mtr4 has been shown to participate in pre-rRNA maturation by directly interacting and recruiting the exosome for processing (de la Cruz *et al*, 1998; Lingaraju *et al*, 2019). Although the phosphosite T34 was detected here, and also previously (Leutert *et al*., 2023), Mtr4 N-terminal 75 amino acids have been shown not to be necessary for the interaction with the exosome (Lingaraju *et al*., 2019). T34 phosphorylation may therefore not be involved in exosome activity control. Mtr4 is a subunit of the TRAMP complex, of which Air2 and Trf4 are also part (LaCava *et al*., 2005). We identified the phosphosite S49 of Air2, in a predicted low complexity region of the protein that controls Mtr4 RNA binding activity (Falk *et al*, 2014). S49 does not directly interact with Mtr4 (distances > 20 Å), but the intrinsically disordered region that contains S49, is enriched in charged residues that promote its high flexibility, suggesting that phosphorylation of S49 might reorient it toward Mtr4. This could stabilize the Air2-Mtr4 interaction and modulate TRAMP complex activity during rRNA processing (Suppl. Fig. 2). Interestingly, although the TRAMP complex localizes to the nucleus, Air2-S49 was only found phosphorylated in the Rrp46 sample, confirming that by using this bait, we isolate both, cytoplasmic and nuclear exosome complexes. These results also further imply a heterogeneity of nuclear exosome complexes, which might have distinct functional roles.

Phosphorylated Ski7, a known cytoplasmic exosome adaptor (Araki *et al*., 2001), was exclusively detected in Rrp46 samples (Fig. 3; Table 1), validating the use of this core subunit as bait for identifying exosome interactors. Phosphosite S90 is present in the middle of the N-terminal portion of Ski7, between the SKI complex-interacting and the exosome-interacting regions of Ski7, whereas S307 is inserted in its GTPase domain (Fig. 6; Suppl Fig. 3), which suggests that the phosphorylation of these residues might affect Ski7 function (Keidel *et al*, 2023; Kowalinski *et al*, 2016).

**Figure 6.**
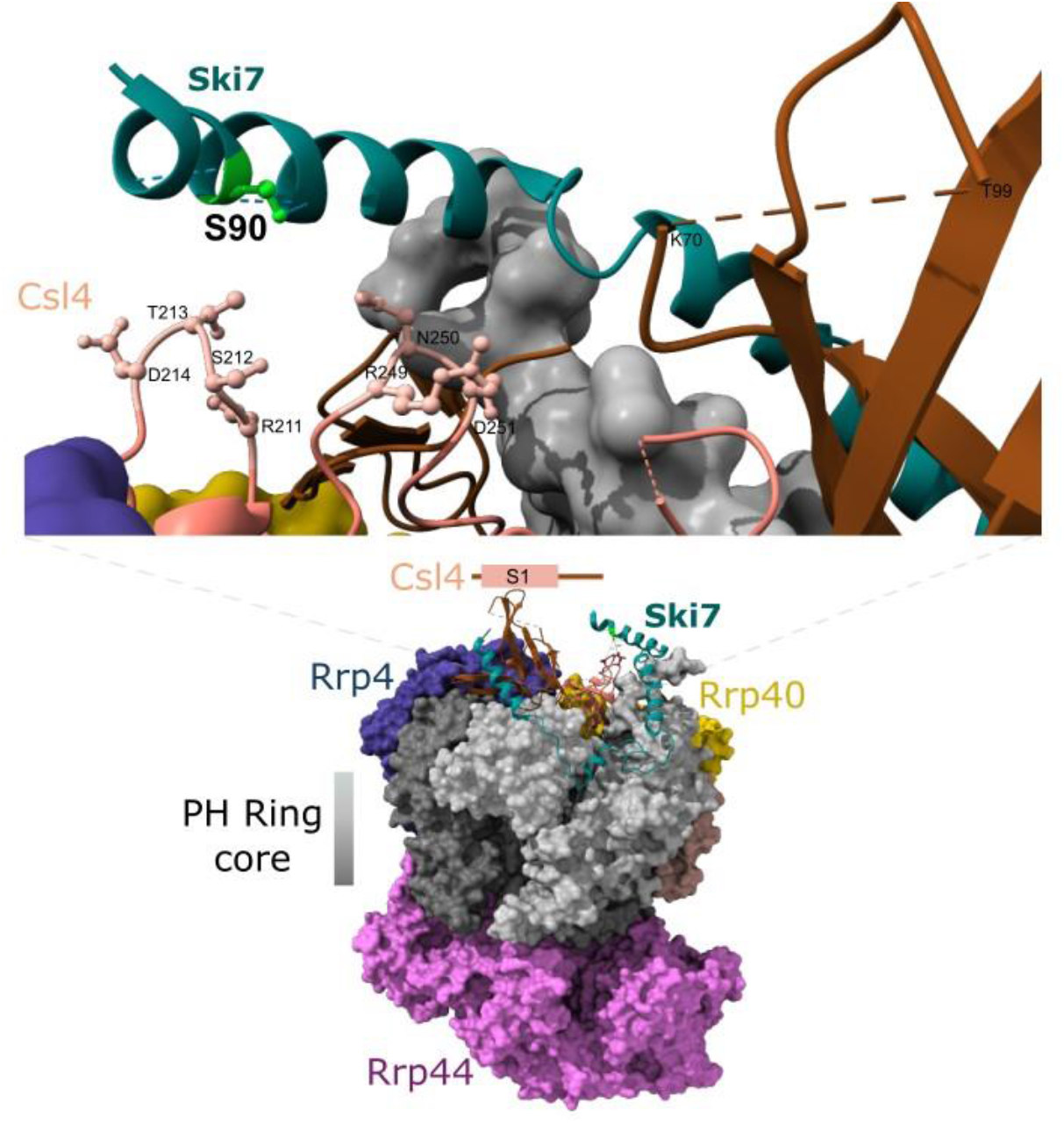
Structure of Ski7 bound to the exosome, highlighting the position of Ski7 phosphorylated residue Ser90 (light green). Structure obtained from PDB 5GO6.

Based on the observation that overexpression of Ski7 traps the exosome in the cytoplasm, therefore affecting ribosome synthesis (Neto *et al*., 2025), to confirm the hypothesis that S90 phosphorylation might have a functional relevance, the genes of wild type Ski7 and the mutant Ski7-S90A were cloned in a plasmid, under control of *MET25* promoter, and transformed into *Δski7* strain to test the effects of their overexpression on cell growth. The results show that while Ski7 overexpression strongly inhibits growth, the non-phosphorylatable Ski7-S90A does not impair growth (Fig. 7), growing very similarly to the condition when Ski7 is expressed at low levels (+Met; Fig. 7). These results confirm the importance of Ski7 phosphorylation for the regulation of its function.

**Figure 7.**
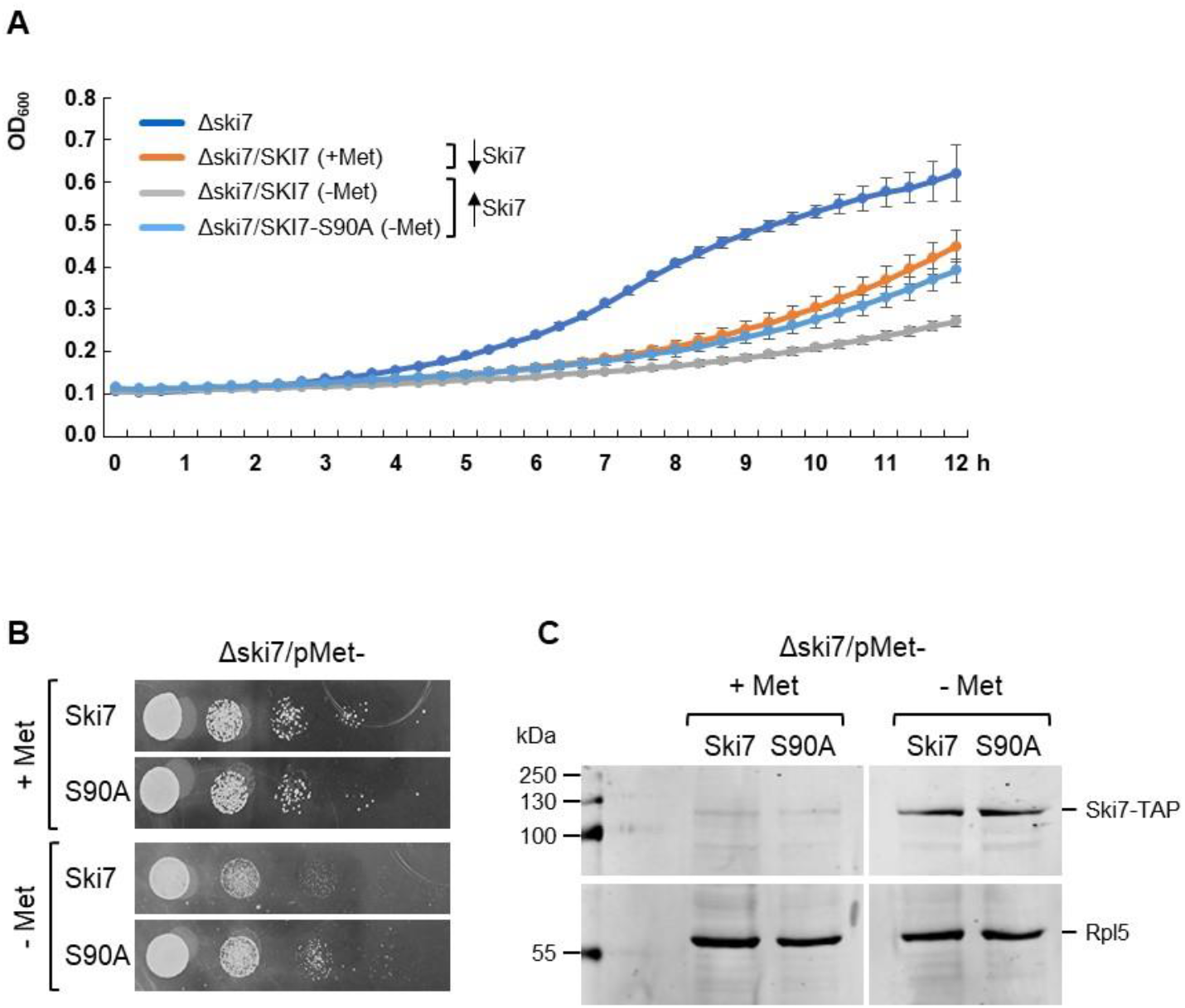
Effects of overexpression of Ski7 on cell growth. Plasmid-borne wild type Ski7, or mutant Ski7-S90A were overexpressed in ¡\ski7 cells and growth was monitored over time. +Met, repression of Δ*Ski7* expression; -Met induction of Ski7 expression. (**A**) Growth curve in liquid medium. (**B**) Serial dilution of cells in OD 0.1 on plates with solid medium. (**C**) Western blot for detection of plasmid-encoded WT and mutant (S90A) Ski7 fused to TAP tag, whose expression is activated in the absence of Met in the medium. Rpl5 was used as loading control.

SSU processome subunit Utp14 phosphosites were also recovered in our samples, with the phosphoresidues S423, S424, and S488 identified both in Rrp6 and Rrp46 eluates (Table 1). These residues are present in Utp14 portion involved in the recruitment of the DEAH-box helicase Dhr1 (Buzovetsky & Klinge, 2025) and, therefore, the phosphorylation of these Ser residues might have functional relevance. Interestingly, we also identified phospho S738 exclusively in Rrp6 samples. This residue is located in the Utp14 region responsible for Dhr1 activation, which occurs right after the release of Rrp6 from the SSU processome (Buzovetsky & Klinge, 2025). Importantly, all the Utp14 phosphosites have previously been identified in global yeast proteomics analyses (Leutert *et al*., 2023), validating our results.

To investigate the evolutionary relevance of the phosphorylation sites identified here, we assessed the conservation of phosphorylated residues across orthologous proteins in human and mouse. The sequence alignments identified conserved phosphorylation sites in corresponding positions of the orthologous sequences. The majority of phosphosites (approximately 58%) were not conserved in either human or mouse orthologs, but a subset of residues (12.4%) was conserved in both mammalian species, including those of Air2 (S49), Ctr9 (S1015, 1017), Enp2 (S550, S555), Nop13 (S2), Pxr1 (S230), Rps21B (S68), Rps6B (S232, S233), Rps7A (T117, T119), Rrp3 (S43, S45, S47), and Rrp4 (S152) (Table 2). The observed conservation of these phosphoresidues suggests potential functional importance across eukaryotes.

**Table 2.**
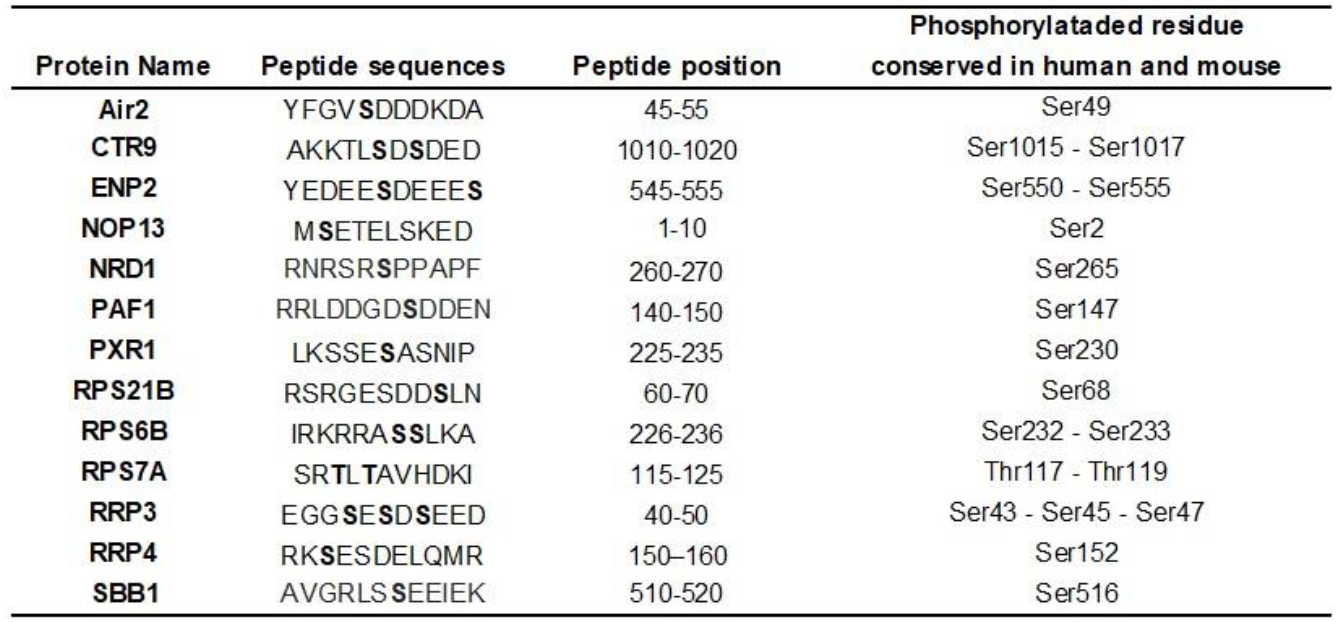
Identified phosphosites that are conserved in human mouse. Partial list of proteins identified, highlighting exosome cofactors, ribosome assembly factors and ribosomal proteins. Phosphoresidues in each protein are indicated.

To evaluate the functional importance of the identified phosphosites, we performed a broad evolutionary analysis, focusing on ribosomal proteins and ribosome assembly factors. We aligned the phosphopeptides identified for these proteins, with their corresponding sequences from 16 additional organisms and observed that many of these conserved peptides also contain conserved phosphoresidues. We then analysed the relative proportions of serine, threonine and tyrosine (STY) residues, as well as aspartic acid and glutamic acid (DE), relative to the total conserved peptides (Fig.8).

The *Saccharomycetaceae* clade has a strong conservation of STY-containing peptides (Fig. 8A, blue bars). As expected, the degree of conservation decreases as more evolutionarily distant the organisms are from *S. cerevisiae*, as indicated by an increase in residues unrelated to phosphorylation (grey bars). In more distant organisms, the conservation of phosphorylatable “STY” residues decreases to approximately 25%. Interestingly, this reduction is accompanied by an increase in phosphomimetic (negatively charged) amino acids (D or E), represented by orange bars. Although these organisms have lost their phosphorylatable residues at these positions, their replacement by phosphomimetic residues suggests that the negative charge at these sites is important for protein structure or function (Fig. 8A).

**Figure 8.**
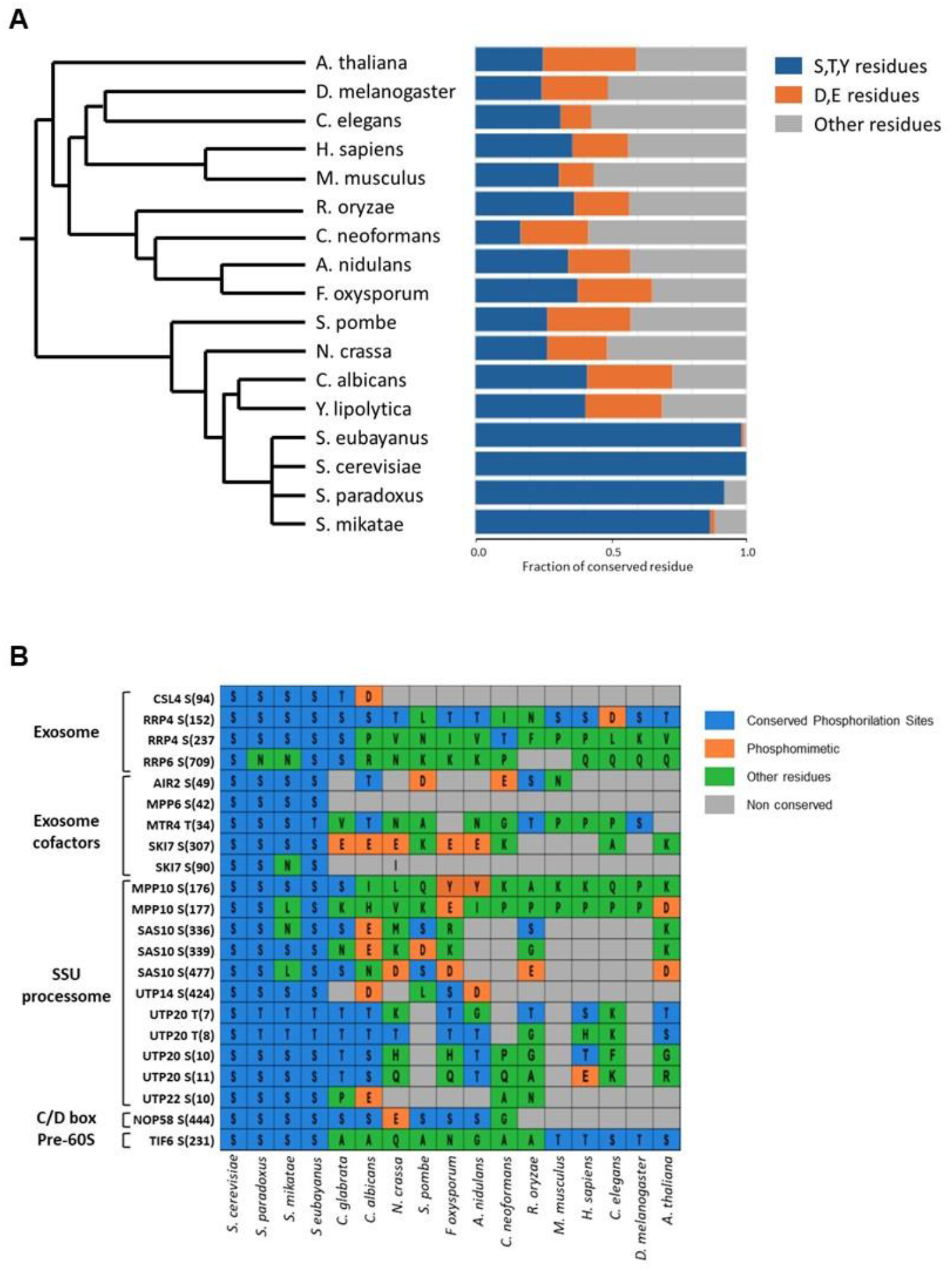
Analysis of phopho peptides conservation. (**A**) Phylogenetic tree showing the percentage of conserved ribosome residues. The tree was constructed using 16 organisms in addition to S. cerevisiae, and the clades were generated using the iTOL (iterative Tree of Life, https://itol.embl.de/itol.cgi) program. Ribosomal associated residues identified in our phosphoproteome analysis were used to generate the alignment and to calculate the relative fraction of conserved residues to each species. (**B**) Plot showing the conservation of phosphoresidues in ribosomal proteins or assembly factors. Ribosomal associated residues identified in our phosphoproteome analysis were used to generate the alignment and to calculate the relative fraction of conserved residues to each species. The analyzed residues were grouped by complex and colored according to their phosphorylation-related characteristics. Blue, “S.T,Y” residues with conserved serine, threonine, or tyrosine. Orange, conserved residues in which STY is replaced by aspartic acid or glutamic acid (;’D, E’’). Green, other residues different from STY or DE. Gray, residues that are not conserved.

The identified *S. cerevisiae* phosphoresidues of exosome subunits and cofactors, and of ribosome assembly factors were aligned with sequences from 16 additional organisms and grouped by complex to analyze the evolutionary conservation of the phosphorylation. The conservation profile mirrors the one presented above, showing the phosphorylatable S, T, Y residues conserved positions in *Saccharomycetaceae*, while D, E substitute these residues in more distant organisms, preserving the negative charge (Fig. 8B). Interestingly, several residues are conserved from yeast to higher eukaryotes. This conservation underscores the importance of these phosphorylation sites for the structure/ function of the RNA exosome complex and ribosome assembly factors, highlighting the role of phosphorylation as a regulatory mechanism in RNA metabolism.

## DISCUSSION

The RNA exosome is involved in processing and degradation of all classes of RNA, both in nucleus and cytoplasm (Chlebowski *et al*, 2013; Januszyk & Lima, 2014; Lykke-Andersen *et al*, 2009; Schneider & Tollervey, 2013). In each cell compartment, the exosome interacts with and is regulated by a specific set of cofactors and interactors (Zinder & Lima, 2017), and these protein interactions may be affected by phosphorylation. In this study, we present a phosphoproteomic investigation of proteins associated with the *Saccharomyces cerevisiae* exosome by using subunits Rrp6 and Rrp46 as baits, and show evidence that phosphorylation may contribute to the structural and regulatory dynamics of the exosome in association with distinct pre-ribosomal intermediate particles and cytoplasmic cofactors.

The majority of proteins interacting with the exosome participate in RNA processing pathway and our data show that these factors exhibit a large number of phosphorylated residues, with a predominance of serine phosphorylation, consistent with global eukaryotic phosphoproteome patterns (Ochoa *et al*, 2020). Phosphorylation was detected both within structural domains and in intrinsically disordered regions (IDRs), the latter being known for facilitating dynamic protein interactions. Our findings are in agreement with previous studies demonstrating that the recruitment of the exosome to pre-ribosomes is mediated by cofactors (Schneider & Tollervey, 2013; Sloan *et al*, 2012) and show their phosphorylation status in distinct complexes, which may influence those interactions.

Although most of the phosphosites isolated here have already been identified in global analyses, which further validates our data, we detected phosphoresidues being co-purified with the exosome as part of pre-ribosomal particles, showing the maturation stage in which their phosphorylation is functionally relevant. In addition, we also detected phosphosites that had not been previously described. The evidence that the exosome is associated with pre-ribosomes is given by the number of proteins identified that participate in various pre-rRNA processing steps. We identified proteins associated with the pre-ribosomal particle 90S (e.g. Utp14, Mpp10), pre-60S (e.g. Nop4, Tif6), proteins involved in nuclear transport of ribosomal particles (e.g. Mtr2, Srm1), subunits of snoRNP particles (e.g. Nop58), ribosomal proteins (e.g. Rps7, Rps14) and nuclear pore complex components (e.g. Nup2, Nup159), which give evidence of the participation of the exosome in multiple steps of ribosomal maturation. The distinct phosphosites identified in the two samples further strengthen this conclusion. Based on these differences, we also hypothesize that the exosome complexes present in the nucleus may not be homogeneous, but rather exist as Exo11 (containing Rrp6) or Exo10 (without Rrp6). This conclusion could also be inferred by previous co-immunoprecipitation of the exosome with Rrp43, in which Rrp6 was not co-purified (Lourenco *et al*., 2013), suggesting a loose association of this subunit with the exosome core.

In addition to ribosome factors, we identified the phosphorylated exosome subunits Csl4, Rrp4 and Rrp6. Interestingly, Rrp4-S237 phosphosite was only identified in Rrp46 samples, strengthening the hypothesis of these phosphorylation events being important for function, in this case, probably in the cytoplasm, or as part of the nuclear Exo10. Furthermore, the cytoplasmic exosome cofactor Ski7 was identified exclusively in Rrp46 samples, validating the purification method. The importance of Ski7-S90 phosphorylation was confirmed here by changing this residue to alanine, which abolishes the negative effect of Ski7 overexpression on cell growth. Interestingly, S90 is part of helix H3 in the SKI complex-interacting portion of Ski7, in close proximity to the exosome-interacting region of Ski7 (positions 116-225; Fig. 6) (Kowalinski *et al*., 2016; Liu *et al*, 2016). It is possible, therefore, that phosphorylation of S90 might influence Ski7 interaction with the SKI complex, and also affect its interaction with the exosome, since Ski7 overexpression has been shown to trap the exosome in the cytoplasm (Neto *et al*., 2025). Our results corroborate the hypothesis that S90 phosphorylation is important for function, and might modulate cytoplasmic RNA degradation pathways.

The analysis of the phosphorylated residues across seventeen organisms ranging from yeast to humans show their evolutionary conservation, further strengthening the validity of our results. Also confirming the importance of the phosphorylation sites identified here, posttranslational modifications of *S. pombe* exosome subunits Dis3 and Rrp6, and cofactor Mtr4, have been shown to be essential for normal cell growth and to fine-tune exosome activity regulation (Telekawa *et al*., 2018).

In conclusion, here we have identified ribosome assembly factors that are phosphorylated, modification that may affect their affinity for binding the nintermediate pre-ribosomal particles, and show evidence that the exosome may not be a homogeneous complex in the nucleus, but rather be present as Exo11 and Exo10 + Rrp6 in this cell compartment, and its subunits may also be subject to phosphorylation for functional regulation.

## Acknowledgements

We are grateful to all members of the Oliveira laboratory for help, reagents, advice and discussion.

## Funding

This work was supported by a grant from Fundação de Amparo à Pesquisa do Estado de São Paulo (FAPESP - 20/00901-1 to C.C.O.). F.A.A., V.G.N., R.B.J., M.R.A.B and B.R.S.Q were supported by FAPESP fellowships (2021/14620-7, 22/00071-4, 2025/06224-5, 2021/14137-4, 2023/06344-5, respectively). LPPC L.P.C. was supported by a CNPq (Conselho Nacional de Desenvolvimento Científico e Tecnológico) fellowship.

## Notes

http://www.ebi.ac.uk/pride

### Competing Interest Statement

The authors have declared no competing interest.

## REFERENCES

Allmang C, Kufel J, Chanfreau G, Mitchell P, Petfalski E, Tollervey D (1999a) Functions of the exosome in rRNA, snoRNA and snRNA synthesis. EMBO J 18: 5399–5410

Allmang C, Petfalski E, Podtelejnikov A, Mann M, Tollervey D, Mitchell P (1999b) The yeast exosome and human PM-Scl are related complexes of 3’ --> 5’ exonucleases. Genes Dev 13: 2148–2158

Araki Y, Takahashi S, Kobayashi T, Kajiho H, Hoshino S, Katada T (2001) Ski7p G protein interacts with the exosome and the Ski complex for 3’-to-5’ mRNA decay in yeast. EMBO J 20: 4684–4693

Bagatelli FFM, de Luna Vitorino FN, da Cunha JPC, Oliveira CC (2021) The ribosome assembly factor Nop53 has a structural role in the formation of nuclear pre-60S intermediates, affecting late maturation events. Nucleic Acids Res 49: 7053–7074

Ballesta JP, Rodriguez-Gabriel MA, Bou G, Briones E, Zambrano R, Remacha M (1999) Phosphorylation of the yeast ribosomal stalk. Functional effects and enzymes involved in the process. FEMS Microbiol Rev 23: 537–550

Bohlen J, Roiuk M, Teleman AA (2021) Phosphorylation of ribosomal protein S6 differentially affects mRNA translation based on ORF length. Nucleic Acids Res 49: 13062–13074

Buzovetsky O, Klinge S (2025) Helicase-mediated mechanism of SSU processome maturation and disassembly. Nature

Cepeda LPP, Bagatelli FFM, Santos RM, Santos MDM, Nogueira FCS, Oliveira CC (2019) The ribosome assembly factor Nop53 controls association of the RNA exosome with pre-60S particles in yeast. J Biol Chem 294: 19365–19380

Cerezo EL, Houles T, Lie O, Sarthou MK, Audoynaud C, Lavoie G, Halladjian M, Cantaloube S, Froment C, Burlet-Schiltz O et al (2021) RIOK2 phosphorylation by RSK promotes synthesis of the human small ribosomal subunit. PLoS Genet 17: e1009583

Chlebowski A, Lubas M, Jensen TH, Dziembowski A (2013) RNA decay machines: the exosome. Biochim Biophys Acta 1829: 552–560

de la Cruz J, Kressler D, Tollervey D, Linder P (1998) Dob1p (Mtr4p) is a putative ATP-dependent RNA helicase required for the 3’ end formation of 5.8S rRNA in Saccharomyces cerevisiae. EMBO J 17: 1128–1140

Falk S, Weir JR, Hentschel J, Reichelt P, Bonneau F, Conti E (2014) The molecular architecture of the TRAMP complex reveals the organization and interplay of its two catalytic activities. Mol Cell 55: 856–867

Filipek K, Blanchet S, Molestak E, Zaciura M, Wu CC, Horbowicz-Drozdzal P, Grela P, Zalewski M, Kmiecik S, Gonzalez-Ibarra A et al (2024) Phosphorylation of P-stalk proteins defines the ribosomal state for interaction with auxiliary protein factors. EMBO Rep 25: 5478–5506

Gonzales-Zubiate FA, Okuda EK, Da Cunha JPC, Oliveira CC (2017) Identification of karyopherins involved in the nuclear import of RNA exosome subunit Rrp6 in Saccharomyces cerevisiae. J Biol Chem 292: 12267–12284

Hartung S, Hopfner KP (2009) Lessons from structural and biochemical studies on the archaeal exosome. Biochem Soc Trans 37: 83–87

Houseley J, LaCava J, Tollervey D (2006) RNA-quality control by the exosome. Nat Rev Mol Cell Biol 7: 529–539

Januszyk K, Lima CD (2014) The eukaryotic RNA exosome. Curr Opin Struct Biol 24: 132–140

Keidel A, Kogel A, Reichelt P, Kowalinski E, Schafer IB, Conti E (2023) Concerted structural rearrangements enable RNA channeling into the cytoplasmic Ski238-Ski7-exosome assembly. Mol Cell 83: 4093–4105 e4097

Kilchert C, Wittmann S, Vasiljeva L (2016) The regulation and functions of the nuclear RNA exosome complex. Nat Rev Mol Cell Biol 17: 227–239

Kowalinski E, Kogel A, Ebert J, Reichelt P, Stegmann E, Habermann B, Conti E (2016) Structure of a Cytoplasmic 11-Subunit RNA Exosome Complex. Mol Cell 63: 125–134

LaCava J, Houseley J, Saveanu C, Petfalski E, Thompson E, Jacquier A, Tollervey D (2005) RNA degradation by the exosome is promoted by a nuclear polyade nylation complex. Cell 121: 713–724

Lau B, Cheng JD, Flemming D, La Venuta G, Berninghausen O, Beckmann R, Hurt E (2021) Structure of the Maturing 90S Pre-ribosome in Association with the RNA Exosome. Molecular Cell 81: 293-+

Leutert M, Barente AS, Fukuda NK, Rodriguez-Mias RA, Villen J (2023) The regulatory landscape of the yeast phosphoproteome. Nat Struct Mol Biol 30: 1761–1773

Lingaraju M, Johnsen D, Schlundt A, Langer LM, Basquin J, Sattler M, Heick Jensen T, Falk S, Conti E (2019) The MTR4 helicase recruits nuclear adaptors of the human RNA exosome using distinct arch-interacting motifs. Nat Commun 10: 3393

Liu JJ, Niu CY, Wu Y, Tan D, Wang Y, Ye MD, Liu Y, Zhao W, Zhou K, Liu QS et al (2016) CryoEM structure of yeast cytoplasmic exosome complex. Cell Res 26: 822–837

Lourenco RF, Leme AF, Oliveira CC (2013) Proteomic analysis of yeast mutant RNA exosome complexes. J Proteome Res 12: 5912–5922

Lykke-Andersen S, Brodersen DE, Jensen TH (2009) Origins and activities of the eukaryotic exosome. J Cell Sci 122: 1487–1494

Makino DL, Conti E (2013) Structure determination of an 11-subunit exosome in complex with RNA by molecular replacement. Acta Crystallogr D Biol Crystallogr 69: 2226–2235

Mitchell P (2010) Rrp47 and the function of the Sas10/C1D domain. Biochem Soc Trans 38: 1088–1092

Mitchell P, Petfalski E, Shevchenko A, Mann M, Tollervey D (1997) The exosome: a conserved eukaryotic RNA processing complex containing multiple 3’-->5’ exoribonucleases. Cell 91: 457–466

Neto VG, Cepeda LPP, Queiroz BRS, Cantaloube S, Leger-Silvestre I, Mangeat T, Albert B, Gadal O, Oliveira CC (2025) New insights into nuclear import and nucleolar localization of yeast RNA exosome subunits. Molecular Biology of the Cell 36

Nielsen PJ, Thomas G, Maller JL (1982) Increased phosphorylation of ribosomal protein S6 during meiotic maturation of Xenopus oocytes. Proc Natl Acad Sci U S A 79: 2937–2941

Ochoa D, Jarnuczak AF, Vieitez C, Gehre M, Soucheray M, Mateus A, Kleefeldt AA, Hill A, Garcia-Alonso L, Stein F et al (2020) The functional landscape of the human phosphoproteome. Nat Biotechnol 38: 365–373

Oliveira CC, Gonzales FA, Zanchin NI (2002) Temperature-sensitive mutants of the exosome subunit Rrp43p show a deficiency in mRNA degradation and no longer interact with the exosome. Nucleic Acids Res 30: 4186–4198

Puig O, Caspary F, Rigaut G, Rutz B, Bouveret E, Bragado-Nilsson E, Wilm M, Seraphin B (2001) The tandem affinity purification (TAP) method: a general procedure of protein complex purification. Methods 24: 218–229

Ramos CR, Oliveira CL, Torriani IL, Oliveira CC (2006) The Pyrococcus exosome complex: structural and functional characterization. J Biol Chem 281: 6751–6759

Schilders G, Raijmakers R, Raats JM, Pruijn GJ (2005) MPP6 is an exosome-associated RNA-binding protein involved in 5.8S rRNA maturation. Nucleic Acids Res 33: 6795–6804

Schneider C, Tollervey D (2013) Threading the barrel of the RNA exosome. Trends Biochem Sci 38: 485–493

Schuch B, Feigenbutz M, Makino DL, Falk S, Basquin C, Mitchell P, Conti E (2014) The exosome-binding factors Rrp6 and Rrp47 form a composite surface for recruiting the Mtr4 helicase. EMBO J 33: 2829–2846

Schuller JM, Falk S, Fromm L, Hurt E, Conti E (2018) Structure of the nuclear exosome captured on a maturing preribosome. Science 360: 219–222

Sloan KE, Schneider C, Watkins NJ (2012) Comparison of the yeast and human nuclear exosome complexes. Biochem Soc Trans 40: 850–855

Synowsky SA, van den Heuvel RH, Mohammed S, Pijnappel PW, Heck AJ (2006) Probing genuine strong interactions and post-translational modifications in the heterogeneous yeast exosome protein complex. Mol Cell Proteomics 5: 1581–1592

Telekawa C, Boisvert FM, Bachand F (2018) Proteomic profiling and functional characterization of post-translational modifications of the fission yeast RNA exosome. Nucleic Acids Res 46: 11169–11183

Thoms M, Thomson E, Bassler J, Gnadig M, Griesel S, Hurt E (2015) The Exosome Is Recruited to RNA Substrates through Specific Adaptor Proteins. Cell 162: 1029–1038

Tomioka M, Shimobayashi M, Kitabatake M, Ohno M, Kozutsumi Y, Oka S, Takematsu H (2018) Ribosomal protein uS7/Rps5 serine-223 in protein kinase-mediated phosphorylation and ribosomal small subunit maturation. Sci Rep 8: 1244

van Hoof A, Staples RR, Baker RE, Parker R (2000) Function of the ski4p (Csl4p) and Ski7p proteins in 3’-to-5’ degradation of mRNA. Mol Cell Biol 20: 8230–8243

Wasmuth EV, Lima CD (2017) The Rrp6 C-terminal domain binds RNA and activates the nuclear RNA exosome. Nucleic Acids Res 45: 846–860

Wasmuth EV, Zinder JC, Zattas D, Das M, Lima CD (2017) Structure and reconstitution of yeast Mpp6-nuclear exosome complexes reveals that Mpp6 stimulates RNA decay and recruits the Mtr4 helicase. Elife 6

Zhang E, Khanna V, Dacheux E, Namane A, Doyen A, Gomard M, Turcotte B, Jacquier A, Fromont-Racine M (2019) A specialised SKI complex assists the cytoplasmic RNA exosome in the absence of direct association with ribosomes. EMBO J 38: e100640

Zinder JC, Lima CD (2017) Targeting RNA for processing or destruction by the eukaryotic RNA exosome and its cofactors. Genes Dev 31: 88–100

