## Supplementary Figures for "Phosphorylation patterns of pre-ribosomal proteins associated with the RNA exosome"

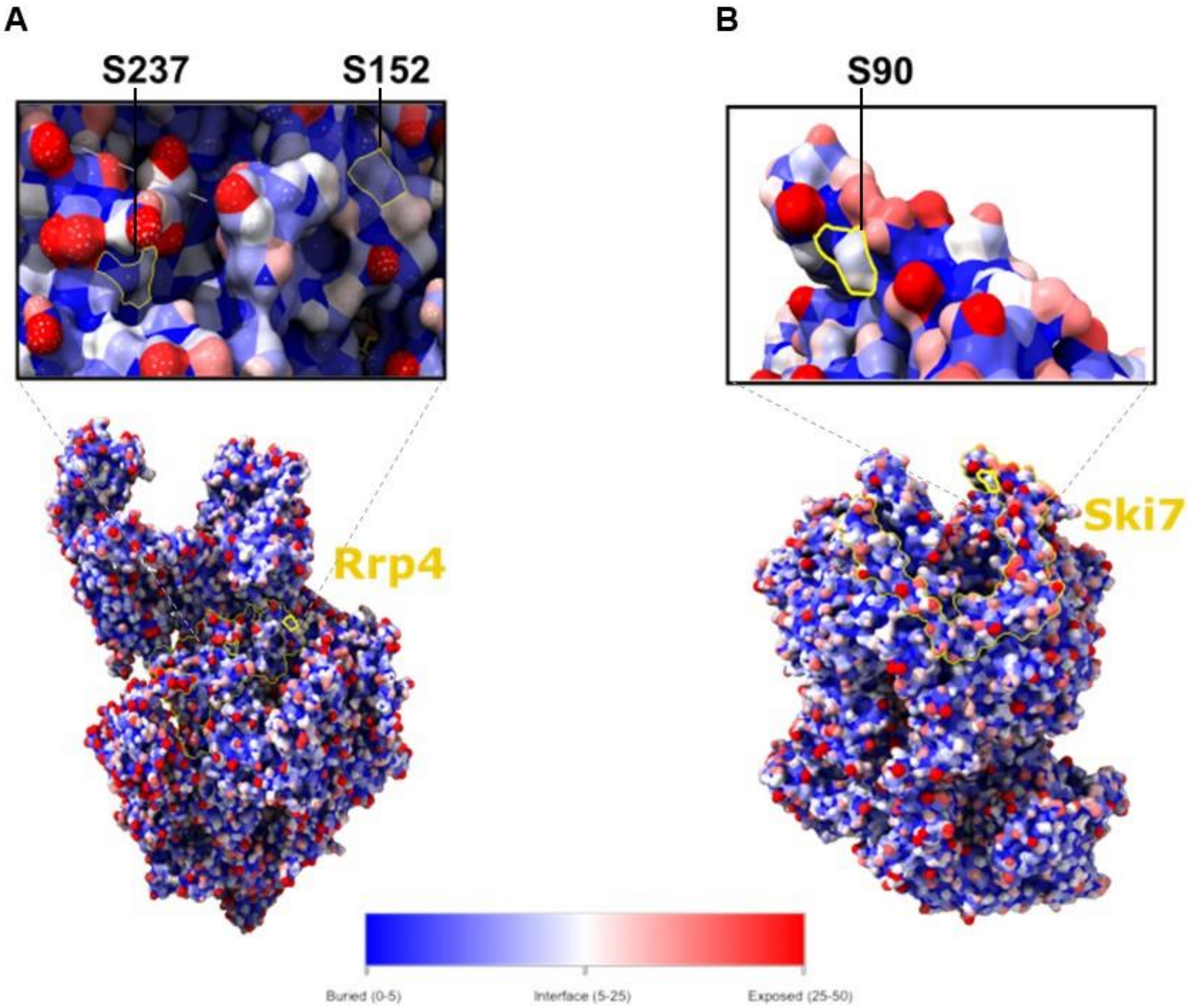


**Suppl. Figure 1. Analysis of the solvent-accessible surface area (SASA) of the *Saccharomyces cerevisiae* exosome complex**. The molecular surface of the *S. cerevisiae* exosome was generated to analyze specific residues identified in this phosphoproteomics study. (**A**) S152 and S237 of Rrp4 (PDB 6FSZ). S152 is located at an accessible interface, while S237 is buried. (**B**) S90 of Ski7 is exposed on the surface (PDB 5G06).

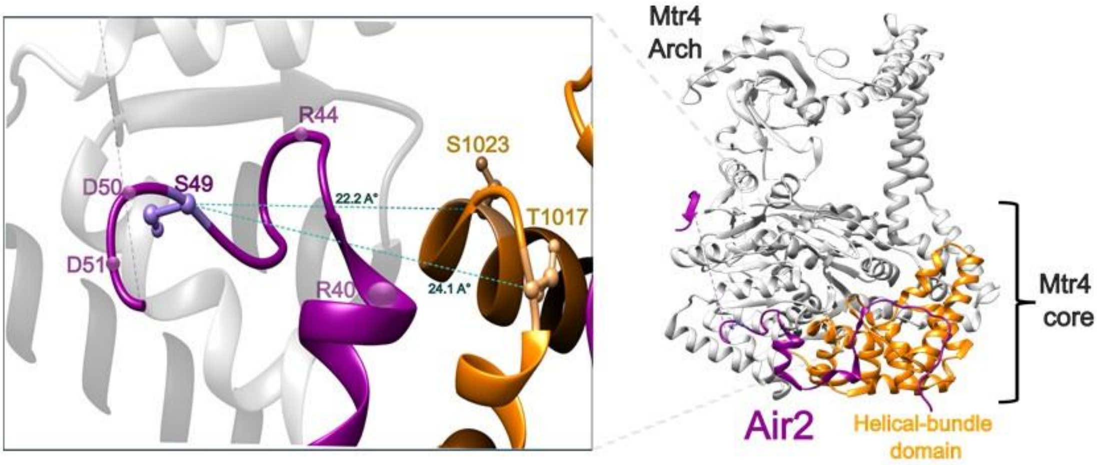


**Suppl. Figure 2.** The phosphorylation site of the Mtr4-Air2 complex. A diagram illustrating the TRAMP complex of *S. cerevisiae* (PDB 4UCU) shows Mtr4 (gray and orange), highlighting the phosphorylation sites. the conserved residues. Air2-S49 (violet) was identified as phosphorylated in this study. It is located in a flexible region flanked by charged residues (R40, R44, D50, D51). Its phosphorylation might activate a conformational switch that establishes electrostatic contacts with the Mtr4 helical bundle domain, coupling the phosphorylation signal to activity in the TRAMP complex during rRNA processing.

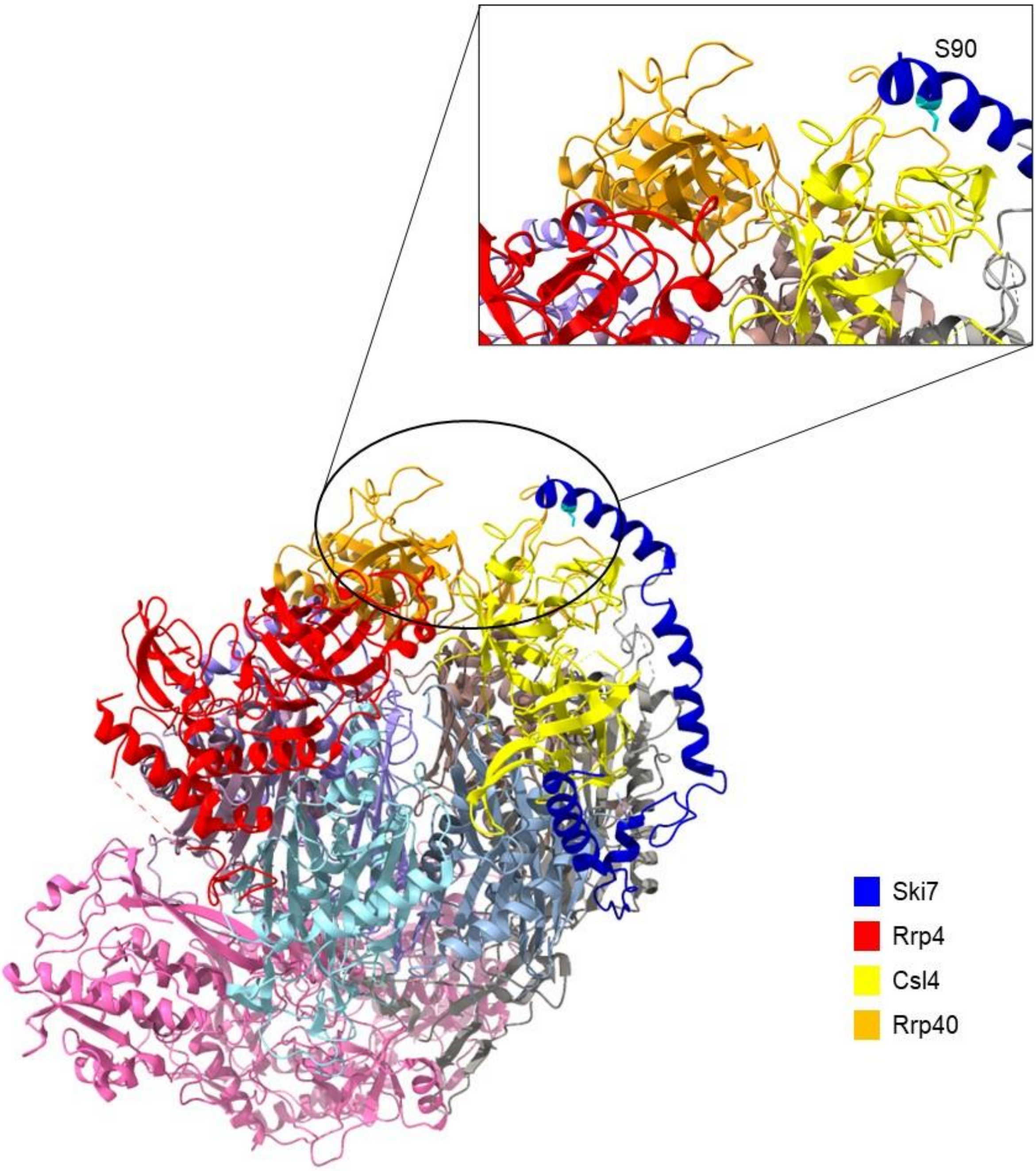


**Suppl. Figure 3.** Ribbon representation of the structure of Ski7 bound to the exosome. The Ski7 phosphorylated residue Ser90 is highlighted in cyan. Structure obtained from PDB 5GO6.
